## Supplement figures 1 to 5 and supplement Tables 1 to 5 for "A Complex Regulatory Landscape Involved In The Development Of External Genitals"

**Short title: *Hox* gene regulation during the development of genitals**

Ana Rita Amândio[1], Lucille Lopez-Delisle[1], Christopher Chase Bolt[1], Bénédicte Mascrez[2] and Denis Duboule[1,2,3,*]

[1]School of Life Sciences, Ecole Polytechnique Fédérale de Lausanne (EPFL), 1015 Lausanne, Switzerland. [2]Department of Genetics and Evolution, University of Geneva, 30 quai Ernest Ansermet, 1211, Geneva, Switzerland. [3]Collège de France, Paris, France.

### Supplementary material

Legends to figures supplements 1 to 5
Figures supplements 1 to 5
Tables supplements 1 to 5

#### LEGENDS TO FIGURES SUPPLEMENTS

**Figure supplement 1: *Hox* genes expression profile during GT development.** Bar plots show the quantification of *Hoxa, Hoxb* and *Hoxc* genes transcripts by RNA-seq (FPKM values) in GT cells at E12.5, E16.5 and E18.5. The gene cluster is indicated on top of each bar plot.

**Figure supplement 2: Quantification of interactions in C-DOM during GT development. A)** Bar plots show the quantification of the ratio of the number of normalized reads (+/- 5Mb around the viewpoint) in selected regulatory regions, using mouse ES cells as a reference. The regulatory element analyzed is indicated on top of each plot. **B)** 4C-seq profiles at the *HoxD* cluster and C-DOM, using GT cells at E12.5, E13.5, E15.5, E17.5 and forebrain cells. Coordinates (mm10): chr2:73815520-74792376. The GT2 (upper panel, blue line) and island V (lower panel, red line) were used as viewpoints.

**Figure supplement 3: Interaction landscape in the *Del(GT2)* allele. A)** ATAC-seq profiles of *HoxD* and C-DOM in wildtype and mutant *Del(GT2)* E13.5 GTs. Coordinates (mm10): chr2:73815520-74792376. The wildtype track is the average of three biological replicates and the *Del(GT2)* track the average of two biological replicates. Peaks called using MACS2 are displayed below, for each individual replicate (vertical black lines below). The red arrows delineate the deleted region and black arrows indicate a peak lost in *Del(GT2)*. **B)** Overlay of 4C-seq profiles of E13.5 GT cells using *Hoxd13* as viewpoint, wildtype in blue and *Del(GT2)* in red (average of two biological replicates). Coordinates (mm10): chr2:73815520-74792376. Viewpoint is highlighted by a gray line. The red arrow indicates the deleted region. **C)** ATAC-seq profile of E10.5 CR (average of two biological replicates). The black arrow points to the GT2 enhancer. Coordinates (mm10): chr2: 73815520-74792376. **D)** RT-qPCR of wildtype and mutant *Del(GT2)* E10.5 CR. *Hoxd13* mRNA levels were analyzed and the values plotted indicate the ratio of expression using wildtype as a reference (blue dots) (n=6 biologically independent WT or mutant GTs). A Welch's *t*-test was used to evaluate the statistical significance of changes in gene expression. Bars indicate mean with SD, * p=0.0125. We observed a 27% decrease in the mRNA levels of *Hoxd13*.

**Figure supplement 4: Chromatin accessibility in the mutant *Del(V)* and *Inv(V)* alleles. A)** ATAC-seq profiles of wildtype and mutant *Del(V)* and *Inv(V)* E13.5 GTs. Coordinates (mm10): chr2:73815520-74792376. The wildtype track is the average of three biological replicates and the *Del(V)* and *Inv(V)* tracks are the average of two biological replicates. Peaks called using MACS2 are displayed below for each individual replicate (vertical black lines below). The red arrows highlight the deleted or inverted region and the black arrow points to the loss of a peak in the *Inv(V)* allele. **B)** Graphical representation of the percentage of interactions centromeric (red) or telomeric (blue) to island V, for each biological replicate. Coordinates (mm10): centromeric: chr2:74015789-74276083; telomeric: chr2:74332870-74671433.

**Figure supplement 5: Alleles generated by CRISPR-Cas9**. Sanger sequencing results of F0 animals for all alleles generated. Scissors indicate CRISPR-Cas9 mediated breakpoints flanking each regulatory region. SgRNA sequences are marked in red or in green. PCR based genotyping was carried out with primers designed on both sides of sgRNAs targets, deletions were screened with primers F1/R2, inversions with primers F1/F2 and R2/R1, and WT were amplified with primers F1/R1 and F2/R2. See Table 1 for all primer sequences and related PCR product sizes.

*Amândio et al.*
Supplementary Figure 1

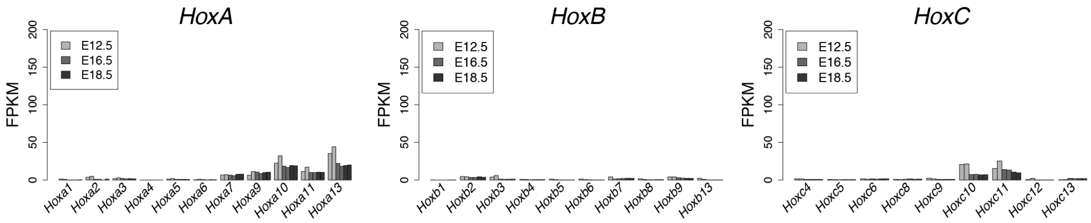

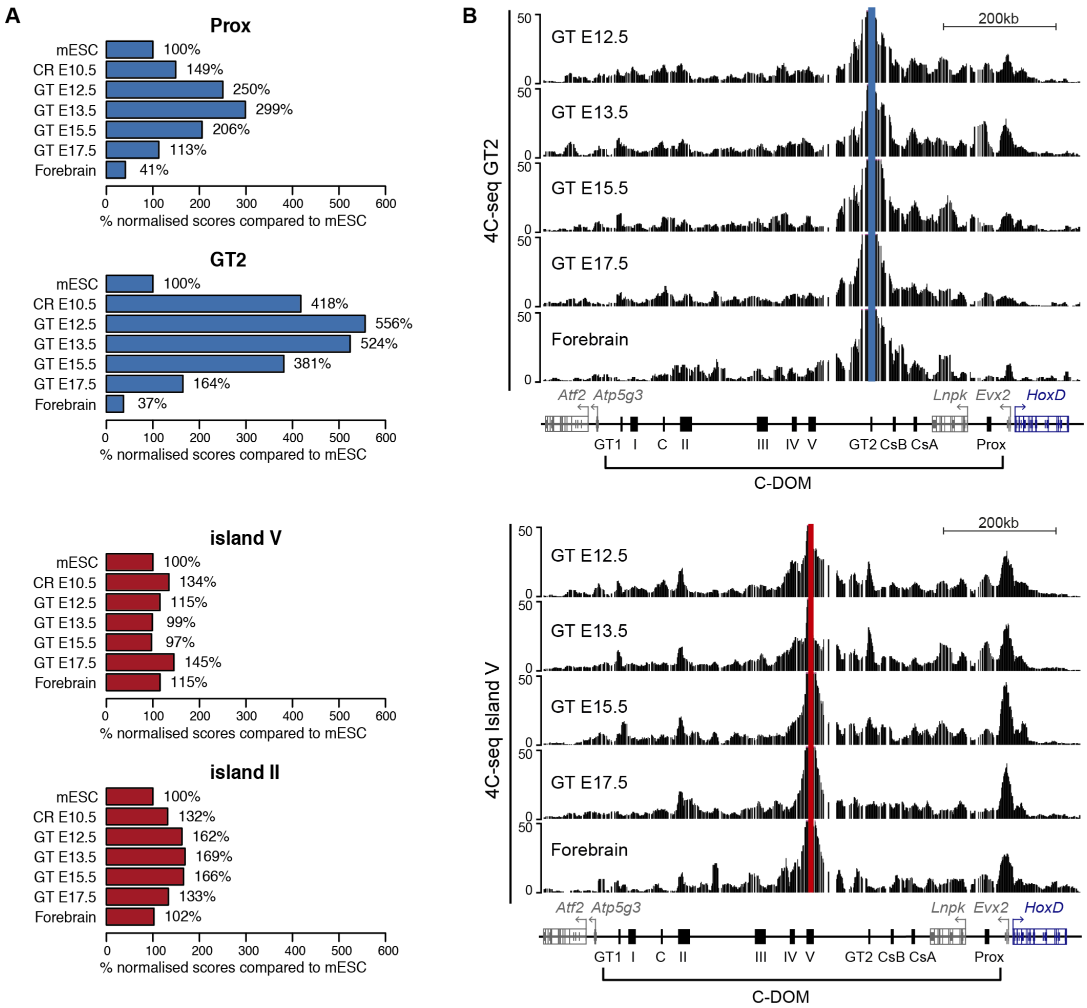
A
Prox
mESC 100%
CR E10.5 149%
GT E12.5 250%
GT E13.5 299%
GT E15.5 206%
GT E17.5 113%
Forebrain 41%
0 100 200 300 400 500 600
% normalised scores compared to mESC
GT2
mESC 100%
CR E10.5 418%
GT E12.5 556%
GT E13.5 524%
GT E15.5 381%
GT E17.5 164%
Forebrain 37%
0 100 200 300 400 500 600
% normalised scores compared to mESC
island V
mESC 100%
CR E10.5 134%
GT E12.5 115%
GT E13.5 99%
GT E15.5 97%
GT E17.5 145%
Forebrain 115%
0 100 200 300 400 500 600
% normalised scores compared to mESC
island II
mESC 100%
CR E10.5 132%
GT E12.5 162%
GT E13.5 169%
GT E15.5 166%
GT E17.5 133%
Forebrain 102%
0 100 200 300 400 500 600
% normalised scores compared to mESC
B
4C-seq GT2
GT E12.5
GT E13.5
GT E15.5
GT E17.5
Forebrain
50
0
200kb
Atf2 Atp5g3 Lnpk Evx2 HoxD
GT1 I C II III IV V GT2 CsB CsA Prox
C-DOM
4C-seq Island V
GT E12.5
GT E13.5
GT E15.5
GT E17.5
Forebrain
50
0
200kb
Atf2 Atp5g3 Lnpk Evx2 HoxD
GT1 I C II III IV V GT2 CsB CsA Prox
C-DOM

*Amândio et al.*
Supplementary Figure 3

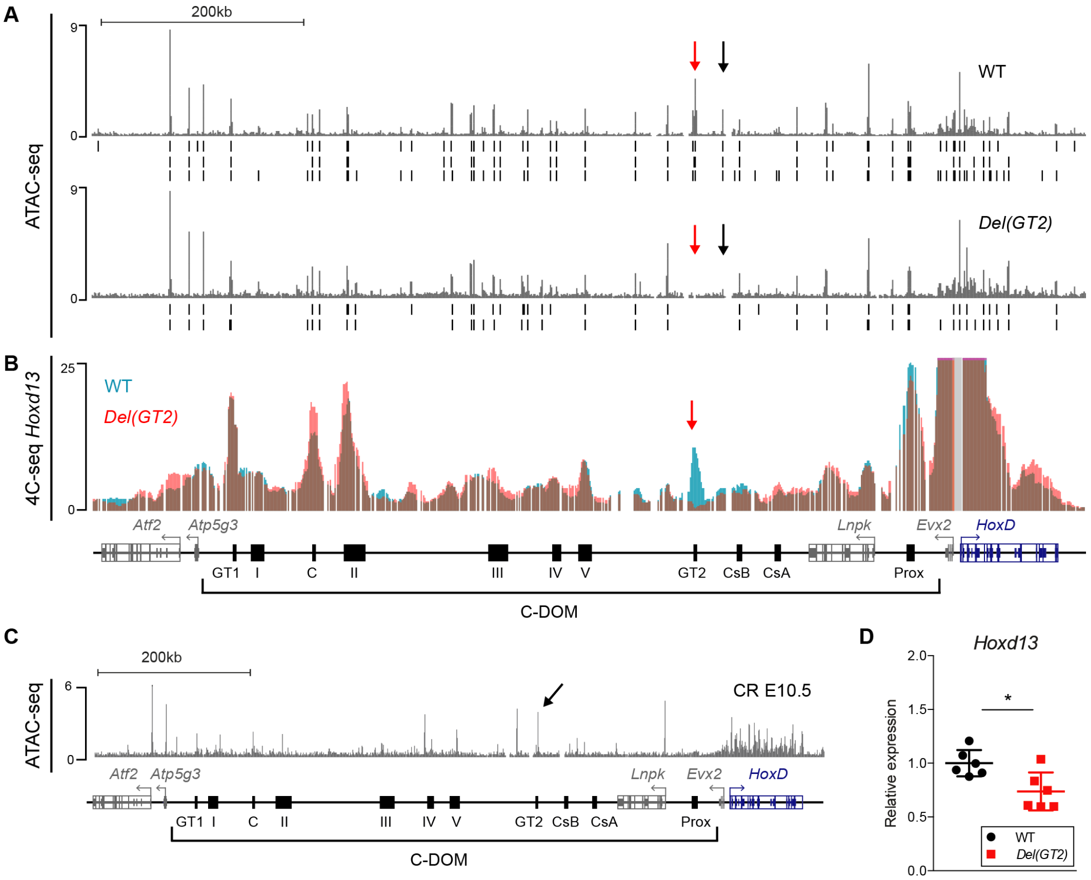

*Amândio et al.*
Supplementary Figure 4

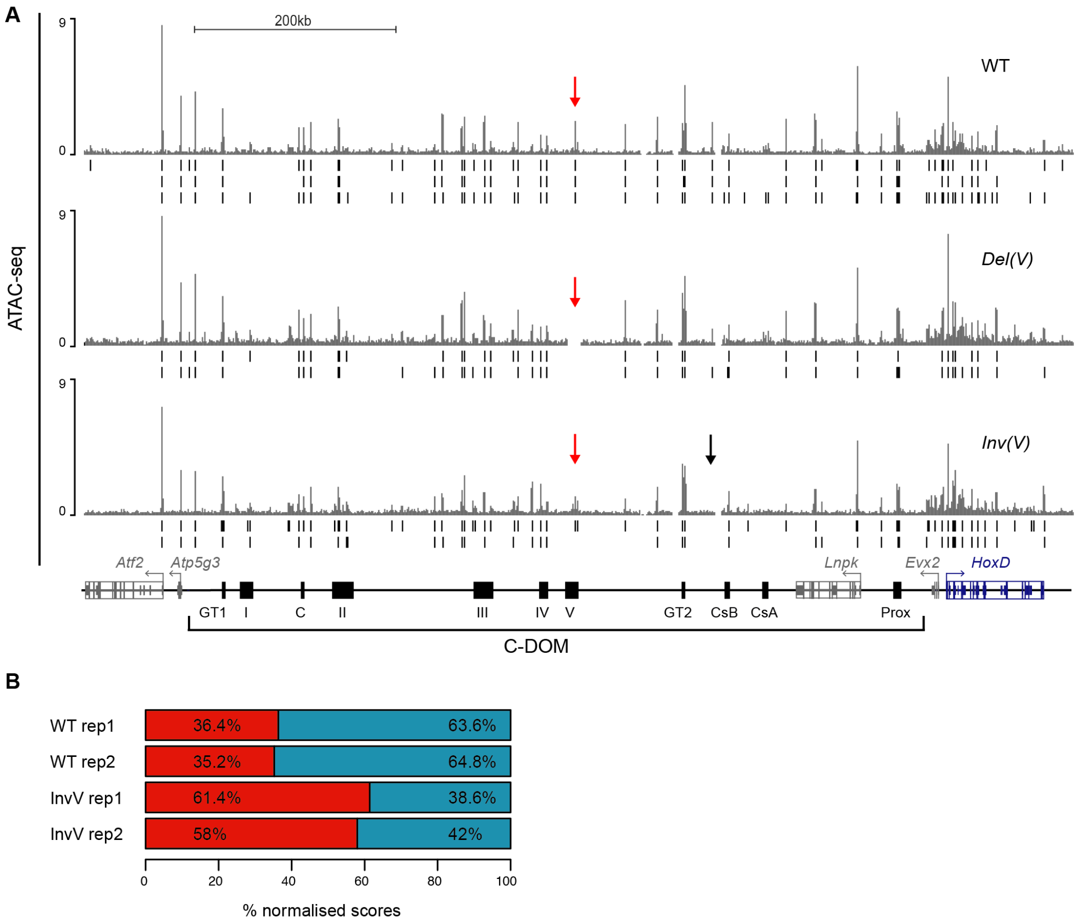

*Amândio et al.*
Supplementary Figure 5

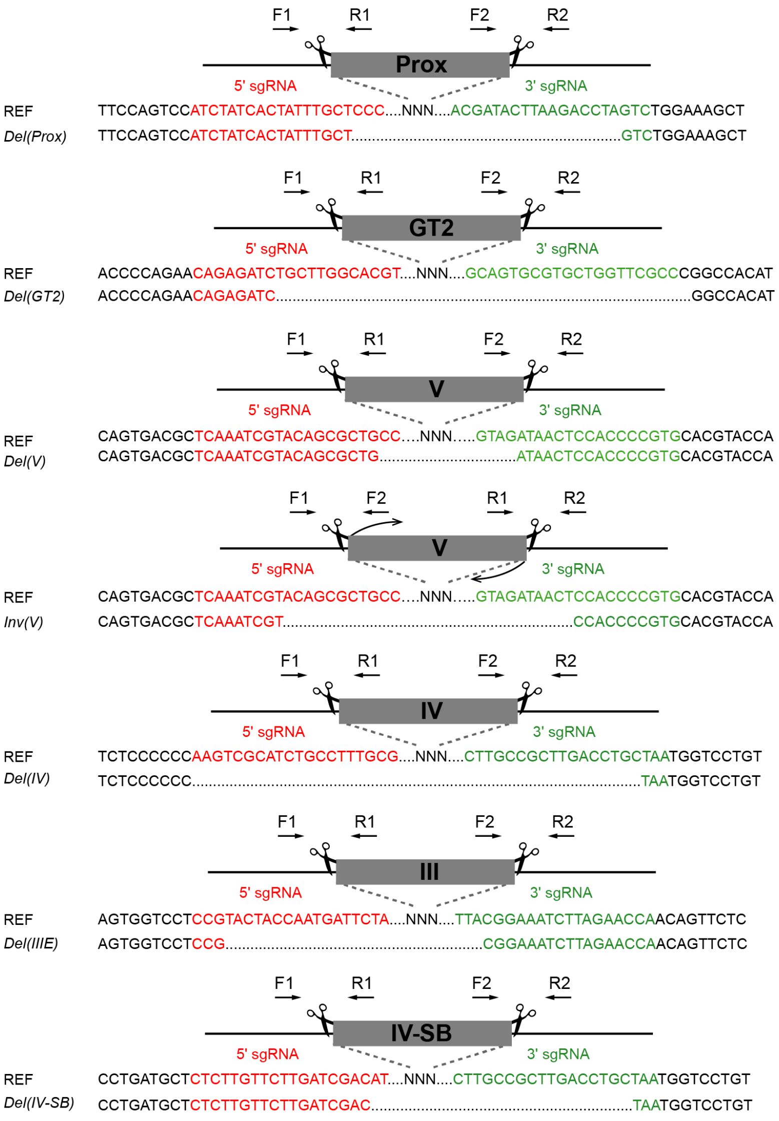

Table 1: List of genotyping primers

| Allele | Primer name | Sequence | Product Size (bp) |
|---|---|---|---|
| *Del(Prox)* | 590 | GACTGTGTTTTGAGGAGGACAGTG | PCR 590-591: WT: 488bp PCR 594-597: Del:369bp |
| | 591 | CTGTGGCAGTAAGAGTGTGCTGAG | |
| | 594 | CCCAGCTGCAGCAGAAGACC | |
| | 597 | GTTTCAAAAGGCAAGTCCATGACTCTCTG | |
| *Del(GT2)* | 435 | TGGGTACAGTTTGGCTCCAT | PCR 435-436: WT: 575bp PCR 433-436: Del: 500bp |
| | 436 | GCCCTTGGTGGCATGTTTAG | |
| | 433 | CTGCCACCTACCTTCTCCTC | |
| *Del(V)* and *Inv(V)* | RA7 | GCTTGTGTCTTGCTGTGTCA | PCR RA7-RA8: WT: 419bp PCR RA7-RA10: Del:440bp PCR RA7-RA9: Inv: approx 300bp |
| | RA8 | AAGGAAAGTGTGTGTGCTGG | |
| | RA9 | CCGAATCCCTAGCTGTCGAG | |
| | RA10 | CATCTGTAGGTTCTGTGCTTATG | |
| *Del(IV)* | 502 | CAGTTCTTTTCCACGGTGAGGAAGC | PCR 502-503: WT: 504bp PCR 501-507: Del: approx 450bp |
| | 503 | CACAAAGATGTGTCAAAGTTGGGGCTG | |
| | 501 | CCTCTTGAACCACGTGCATCGC | |
| | 507 | GCCAGCAGGAATAGCTATACTAACAGG | |
| *Del(III)* | 545 | GGAGAGCTCTGGGTGTGATTGC | PCR 545-542: Wt: 397bp PCR 537-544: Del: 600bp |
| | 542 | TGGGAAGTGTGGAGTCTTCTGCC | |
| | 537 | CCCTTCCCCCTATCACTGTATCTCC | |
| | 544 | CAGACCTTTGCAGTAGGGTCATGG | |
| *Del(IV-SB)* | RA21 | TCTGCCTCCGTTCTCACAAT | PCR RA21-RA22: Wt: 480bp PCR RA21-RA24: Del:300bp |
| | RA22 | GGACCATCAAGAAGCATCCG | |
| | RA24 | GCACTAATCCAAAGCCAGCA | |

Table 2: List of sgRNAs

| **Allele** | **5' sgRNA sequence** | **3' sgRNA sequence** |
|---|---|---|
| *Del(Prox)* | ATCTATCACTATTTGCTCCC | ACGATACTTAAGACCTAGTC |
| *Del(GT2)* | CAGAGATCTGCTTGGCACGT | GCAGTGCGTGCTGGTTCGCC |
| *Del(V)/Inv(V)* | TCAAATCGTACAGCGCTGCC | CACGGGGTGGAGTTATCTAC |
| *Del(IV)* | CGCAAAGGCAGATGCGACTT | CTTGCCGCTTGACCTGCTAA |
| *Del(III)* | TAGAATCATTGGTAGTACGG | TGGTTCTAAGATTTCCGTAA |
| *Del(IV-SB)* | CTCTTGTTCTTGATCGACAT | CTTGCCGCTTGACCTGCTAA |

Table 3: List of fosmids

| Clone Name | Coordinates | Length |
|---|---|---|
| WI1-1879J12 (island IV) | chr2:74235922-74281101 | 45180 bps |
| WI1-2556D5 (island III) | chr2:74180517-74223072 | 42556 bps |
| WI1-1741G6 (island V) | chr2:74270669-74309212 | 38544 bps |
| WI1-1129A16 (between is-III/SB) | chr2:74141402-74180933 | 39532 bps |
| WI1-109B16 (between is-V/GT2) | chr2:74346499-74384278 | 37780 bps |

Table 4: List of primers used for recombineering

| **Primer name** | **Sequence 5' > 3'** |
|---|---|
| WI1-2556D5 (island III) Fw | **CAGTTCTCAGTCTTCAACTTGCTGAGTCAAAAATCTGTTGTTCCTGTATT**GCTCGAGGTCGACGGT ATCG |
| WI1-2556D5 (island III) Rev | **CTTTTCTGTGCTATTCTGAGGAGTGTTGGTGTTGTTACTTCTGGGAAGTG**ATCTATGTCGGGTGCGG AGAAAGAGGTAATGAAATGG CGTCCGCCATCTCCAGCAGC |
| WI1-1879J12 (island IV) Fw | **AATAAATGAACAGCGCTGCTCAGCTGCCCCTTCCCCCGTGGCTAGGCATT** GCTCGAGGTCGACGGTATCG |
| WI1-1879J12 (island IV) Rev | **TCTTAACAGAGTAGCTTCCTATGTGGAATGTCTGTGGTGGAAAGCGAGGA**ATCTATGTCGGGTGCG GAGAAAGAGGTAATGAAATGG CGTCCGCCATCTCCAGCAGC |
| WI1-1741G6 (island V) Fw | **ACTGTTCCCTAATTAATACTAAGCCATATGCAAAAACATTCTTGAATACC**ATCTATGTCGGGTGCG GAGAAAGAGGTAATGAAATGGGCTCGAGGTCGACGGTATCG |
| WI1-1741G6 (island V) Rev | **TCTTAACAGAGTAGCTTCCTATGTGGAATGTCTGTGGTGGAAAGCGAGGA** CGTCCGCCATCTCCAGCAGC |
| WI1-1129A16 (between is-III/SB) Fw | TTTTGATCCTGAGGCAGCGGAAGACTGTGCCACACTAGATGTAACTTGAG GCTCGAGGTCGACGGTATCG |
| WI1-1129A16 (between is-III/SB) Rev | TGATTCTTACCAGACTCTAGTTGTCTGCATCCCAAGTTCCCTACTAGGAC ATCTATGTCGGGTGCGGAGAAAGAGGTAATGAAATGG CGTCCGCCATCTCCAGCAGC |
| WI1-109B16 (between is-V/GT2) Fw | **CACCCAGTTTTTTAATCCTGCCGGATCTTTGCATTGACGGTGGCTGTAA**GCTCGAGGTCGACGGT ATCG |
| WI1-109B16 (between is-V/GT2) Rev | **CAAGCATCTCCTCTTTGGTGTCTCAGAAGCAGTTGGCTAAATGAGCTCCT**ATCTATGTCGGGTGCG GAGAAAGAGGTAATGAAATGG CGTCCGCCATCTCCAGCAGC |

**Homology Arm** - restriction site (PISceI) - Primer

Table 5: List of 4C-seq primers

| Viewpoint | | Sequence (5' > 3') |
|---|---|---|
| *Hoxd13* | iF | AATGATACGGCGACCACCGAACACTCTTTCCCTACACGACGCTCTTCCGATCT<br>XXXXAAAATCCTAGACCTGGTCATG |
| | iR | CAAGCAGAAGACGGCATACGAGGCCGATGGTGCTGTATAGG |
| GT2 | iF | AATGATACGGCGACCACCGAACACTCTTTCCCTACACGACGCTCTTCCGATCT<br>XXXXTTCTCTCTTTTAGTGACCTTGGAACA |
| | iR | CAAGCAGAAGACGGCATACGAAGAAATATCCAAAGGTAAAAATCAAGAA |
| island V | iF | AATGATACGGCGACCACCGAACACTCTTTCCCTACACGACGCTCTTCCGATCT<br>XXXXGCTACAAGACTCATTCGTTTAA |
| | iR | CAAGCAGAAGACGGCATACGAAACTAACTTAAGTCCCCTCG |
| island IV | iF | AATGATACGGCGACCACCGAACACTCTTTCCCTACACGACGCTCTTCCGATCT<br>XXXXTACAGCCTAGTCTTTTCTCATCACAT |
| | iR | CAAGCAGAAGACGGCATACGATGTAATTATTTCAGGGTTGGAGTAGAATCA |

XXXX – Corresponds to possible barcode sequences
